# Spatially distinct chromatin compaction states predict neoadjuvant chemotherapy resistance in Triple Negative Breast Cancer

**DOI:** 10.64898/2025.12.04.692131

**Authors:** Reut Mealem, Thomas. A. Phillips, Leor Ariel Rose, Stefania Marcotti, Maddy Parsons, Assaf Zaritsky

**Author notes:** Co-corresponding authorship Assaf Zaritsky Maddy Parsons. Equal contribution (Reut Mealem, Thomas. A. Phillips, Leor Ariel Rose).

## Abstract

Organisation and dynamics of chromatin play a key role in regulation of cell state and function. In cancer, chromatin plasticity is known to be important in control of drug resistance, but the relationship between chromatin compaction and chemotherapy response within complex tissue settings remains unclear. Here, we measured single nuclei chromatin compaction using fluorescence lifetime imaging microscopy (FLIM) in situ in whole biopsies from 53 pre treatment and 14 post surgery tissue samples of human triple-negative breast cancer (TNBC) patients to determine whether single nuclei spatial chromatin compaction state can predict resistance to neoadjuvant chemotherapy (NACT). Bulk chromatin compaction across 53 pre-treatment core biopsies did not predict patient outcome. However, machine learning analysis revealed that a subset of patients exhibited distinct distributions of single nuclei with more open chromatin states, which was predictive of NACT-resistance.

Graph neural network analysis established that the spatial arrangement of chromatin compaction contributed to prediction of NACT resistance and that chromatin compaction signatures were preserved in tissue state transitions from pre- to 14 post-NACT samples. Our findings shed new light on spatial control of chromatin structure and relationship to therapeutic resistance and establish a foundation for further molecular analysis of chromatin states in complex biological tissues.

## Introduction

The intricate and dynamic 3D organisation of chromatin is critical to control of genome function. Different degrees of chromatin condensation regions coexist within single nuclei, and this organization is continuously shaped by chromatin regulators and in response to cues from the microenvironment ^1,2^. Chromatin is involved in many essential functions such as DNA replication and repair ^3^, cell-cycle progression ^4,5^, programmed cell death ^6^, cell differentiation and transcriptional regulation ^7^. Alterations to chromatin structure, for example via DNA methylation, histone modifications, or chromatin remodeling, has been linked to a broad spectrum of diseases ^8,9^, including cancer ^10–16^. High-dimensional features of chromatin structure extracted from single cell images can distinguish between different histological annotations ^17,18^, between different cancer cell states in patient derived tissue biopsies^17^ and between different disease stages ^18^. Changes in chromatin organisation have also been linked to different breast cancer subtypes ^19^, where more aggressive triple-negative breast cancer (TNBC) cells exhibit dramatic alterations in 3D chromatin structure, with highly disorganized DNA folding, compared to other breast cancer subtypes ^20^.

Chromatin plasticity is particularly evident in cancer cells that develop chemotherapy resistance: these cells can transiently compact their chromatin immediately after treatment followed by decondensation during recovery ^21^. However, the nature of chromatin compaction in chemotherapy-resistance remains controversial and context-specific.

Chemotherapy-resistant cancer cells can adopt more compact chromatin states, which physically protects the DNA, rendering it less accessible to commonly used DNA-intercalating chemotherapies ^22^. Chemotherapy resistance can also be promoted by specific compaction of intrinsically tightly packed heterochromatin, contributing to the silencing of DNA repair genes and thus maintaining cell viability under conditions that cause DNA damage ^23^. Conversely, emerging evidence suggests that specific regions of open chromatin can promote resistance to targeted therapies such as PARP inhibitors ^24^.

Chemotherapy resistance can also be supported by specific chromatin re-organization that can enable transcriptional flexibility and upregulation of pro-survival genes ^21^. Moreover, chemical agents that promote chromatin compaction lead to silencing of drug-resistance genes, resulting in greater sensitivity to cisplatin-based chemotherapies despite a more compact chromatin state ^25^. Furthermore, most previous studies have focused on chromatin compaction in cancer cells, and fail to take into account the diversity of other cell types present within the tumour microenvironment that could contribute to drug resistance ^26,27^.

Here we analysed formalin-fixed paraffin-embedded (FFPE) tissue samples from a large cohort of TNBC patients to determine whether single nuclei level chromatin compaction state can predict resistance to neoadjuvant chemotherapy (NACT). Reducing the chromatin complexity of a single nuclei to a single scalar approximating the chromatin compaction via fluorescence lifetime imaging microscopy (FLIM) ^28^, revealed that nuclei from TNBC patients resistant to NACT exhibited more open chromatin compared to nuclei from responder patients. Machine learning analysis revealed that encoding heterogeneity within the tissue was necessary for the prediction of NACT resistance and that the spatial arrangement of nuclei further contributed to this predictive power. Moreover, the signature of chromatin compaction states within the tissue was preserved from pre-treatment to post-treatment samples in resistant patients, indicating spatially encoded chromatin state may define and perpetuate drug resistance.

## Results

### The distribution of single nuclei chromatin compaction in TNBC patients predicts resistance to NACT

To examine whether the level of chromatin compaction within single nuclei correlates with resistance of TNBC patients to NACT, a retrospective cohort of 53 pre-treatment tissue samples (core biopsies termed “cores”) was assembled for analysis (Fig. 1A). Each patient was classified as “responder” or “non-responder” to NACT based on the residual cancer burden (RCB) score, a measure for the amount of tumor tissue remaining after the NACT treatment period reflecting effectiveness in reducing primary tumor size (Fig. 1B). Responders had RCB scores of 0 or 1, whereas non-responders had RCB scores of 2 or 3. Post-surgery samples (taken at surgery and termed “resections”) were also acquired for “non-responder” patients. The cohort consisted of 30 responder and 23 non-responder cores, and resections for 14 of the non-responders. All tissue samples were stained using SiR-DNA and imaged using FLIM, yielding both a nuclear fluorescence intensity image and a fluorescence lifetime image at single nuclei resolution (Fig. 1C, left). The intensity image reflects the total fluorescence emitted from the dye-labelled nuclei. The lifetime image is independent of dye concentration, and represents the average time the fluorophore remains in its excited state before relaxing to its ground state by emitting a photon. This captures differences in chromatin compaction that arise from changes in the fluorophore local environment affecting its relaxation dynamics ^28,29^. The fluorescence lifetime ranged from 0 to 13 nanoseconds, where higher values denote less condensed and lower values more condensed chromatin^30^. SiR-DNA has been used previously to report on chromatin state; however we also undertook control experiments to ensure the dye was reporting on this organisational feature in our hands. HeLa cells were left untreated, or treated with increasing doses (2, 10, 20%) of dextrose to induce osmotic shock and chromatin compaction ^30^. Cells were then stained with SiR-DNA and imaged using FLIM. Data demonstrated a significant reduction in fluorescence lifetime with increasing doses of dextrose indicating shorter lifetime values correspond to greater degrees of chromatin compaction (Fig. S1), confirming previous reports and probe sensitivity to chromatin organisation.

**Figure 1.**
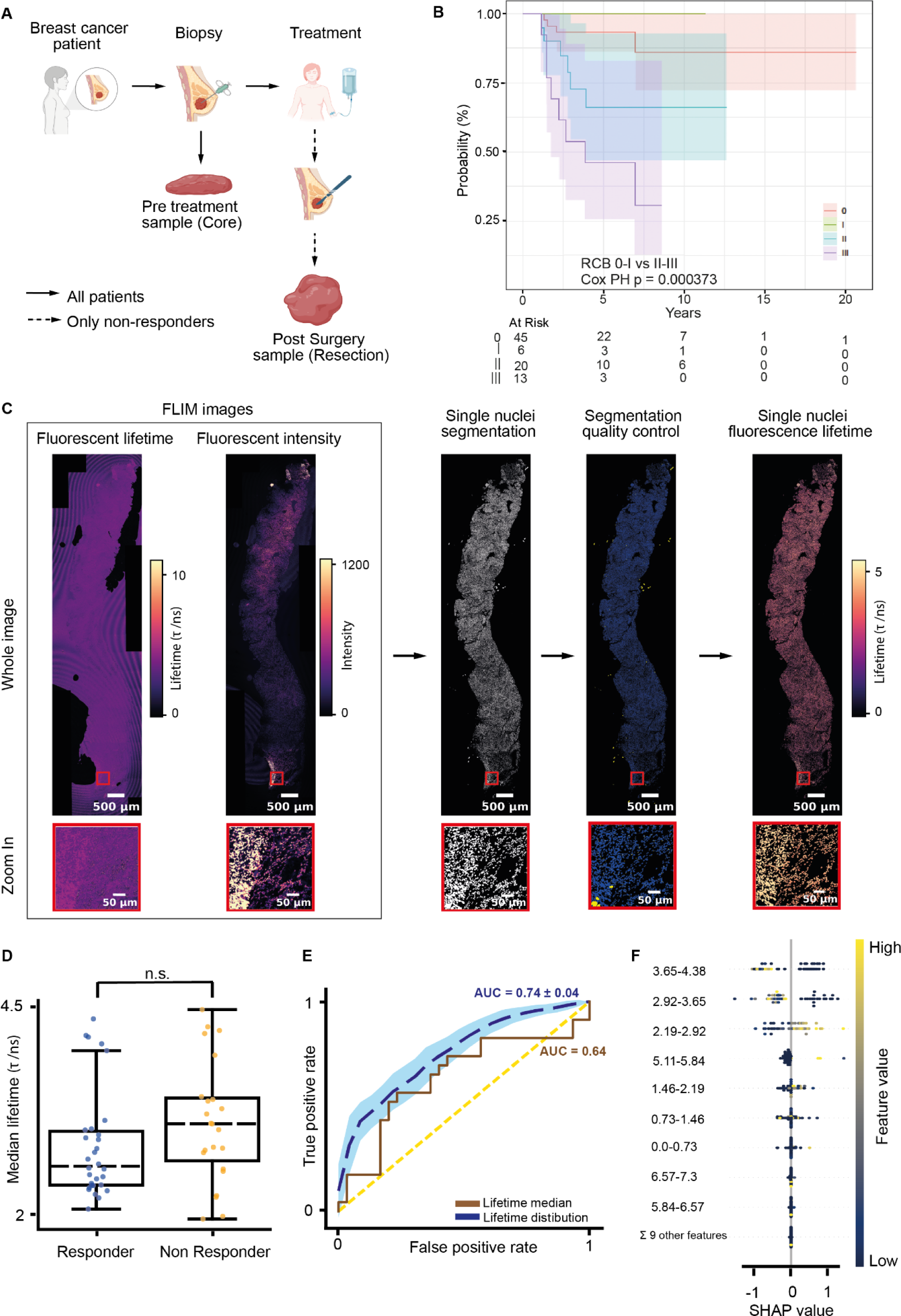
Cohort, analysis pipeline and machine learning prediction of patient treatment response. **(A)** Retrospective TNBC cohort treated with neoadjuvant chemotherapy (NACT). Pre-treatment biopsies (cores) were collected from all patients. Subsequently, all patients received treatment. Post-surgery samples (resections) were acquired for patients who were resistant to chemotherapy (non-responders). **(B)** Kaplan Meier survival plot stratified by Residual Cancer Burden (RCB) classification score. Patients were categorized as responders or non-responders to NACT based on their RCB score, which reflects the amount of residual tumor tissue following treatment. Poor survival outcomes in patients with higher RCB scores (RCB-II and RCB-III) compared to those with minimal or no residual disease (RCB 0 and RCB-I) (p < 0.001, log-rank test. Shaded areas around each survival curve represent 95% confidence intervals. The “At Risk” table shows the number of participants at each time point on the x-axis. **(C)** Image processing workflow. SiR-DNA stained samples were imaged using fluorescence lifetime imaging microscopy (FLIM), yielding fluorescence intensity and fluorescence lifetime images. Nuclei were segmented from the intensity image, manual and automated quality control was applied to exclude segmentation errors (yellow objects, best observed in the zoom-in image). Each nuclei was assigned with its average fluorescence lifetime in its corresponding segmented mask. Red boxes provide zoom-in views at the bottom. Scale bar = 500 𝜇m (whole image), 50𝜇m (zoom in). **(D)** The distribution of responder versus non-responder cores, where each data point represents the median fluorescence lifetime from all single nuclei in a core. Two-sided independent t-test did not reject the null hypothesis (H₀: responder median = non-responder median, p-value = 0.155). **(E)** Receiver Operating Characteristic (ROC) curve for fluorescence lifetime-based cores classification. The average ROC curve of the lifetime distribution-based XGBoost model (blue dashed line, ROC-AUC = 0.74) with the standard deviation (shade) calculated across 100 different random seeds surpassed the ROC curve of the median fluorescence lifetime of each core (brown solid line, ROC-AUC = 0.64). The yellow dashed line Y = X represents random discrimination (ROC-AUC = 0.5). **(F)** SHapley Additive exPlanations (SHAP) analysis for the top-ranked features in the XGBoost fluorescence lifetime distribution model. Each horizontal feature (e.g., “3.65-4.38”) represents a bin in the fluorescence lifetime distribution, with the range indicating the lifetime interval (in nanoseconds). Positive SHAP values indicate features contributing to responder predictions, while negative SHAP values correspond to features contributing to non-responder predictions. Each data point represents SHAP value from one tissue with the color encoding the feature values, i.e., the normalized frequency of nuclei in each bin.

Having validated the probe, the tissue SiR-DNA FLIM images were processed to single nuclei fluorescence lifetime values using the following image analysis pipeline (Fig. 1C, left-to-right). Firstly, individual nuclei in the FLIM fluorescence intensity image were segmented with StarDist ^31^. Secondly, visual quality control excluded segmentation errors from downstream analysis. Finally, the fluorescence lifetime of each nucleus was computed using the fluorescence lifetime image as the average intensity in its segmented mask. This data generation and analysis pipeline represented each tissue sample as the spatial arrangement of single nuclei fluorescence lifetime values. To explore the association between the fluorescence lifetime and nuclei shape or local density, we quantified the eccentricity, size and local density for all nuclei in each tissue sample and found marginal correlation between each of these measurements and the fluorescence lifetime (Fig. S2A).

The median fluorescence lifetime values of single nuclei from cores did not show a statistical difference between responders and non-responders (Fig. 1D), and achieved a marginal ROC-AUC score of 0.64 in distinguishing between these patient groups (Fig. 1E). The cores’ median nuclei eccentricity, size or local density also did not show statistical differences between these patient groups (Fig. S2B-D). This indicates that a single-value summary does not adequately capture fluorescence lifetime variations between the response groups. Extending the core representation from the median fluorescence lifetime to the full distribution of the single nuclei fluorescence lifetime offered a richer depiction that captured the inherent nuclear heterogeneity within the tissue. We then applied machine learning approaches to determine the association between the distribution of fluorescence lifetime to NACT treatment response. Due to the limited number of patients in the cohort, we trained XGBoost ^32^ classification models to predict treatment response with a leave-one-core-out cross validation: multiple rounds of training and testing, where in each round, one core was designated as “test”, and all other cores were used for training. To ensure robustness and account for stochastic variability, we repeated model training (and then testing) across 100 different random seeds, providing a comprehensive assessment of performance stability. Classification performance achieved a median ROC-AUC of 0.735 (maximum of 0.850) (Fig. 1E). To confirm that these results were not due to overfitting, we shuffled the NACT response labels, retrained the classification models and recorded the shuffled ROC-AUC. Repeating this process 1000 times and assessing how many different shufflings induced performance equal or better than the median ROC-AUC observed with the true (i.e., non-shuffled) labels yielded a p-value of 0.004, establishing that the observed classification was not due to chance. To pinpoint which interval ranges of fluorescence lifetime values were the most influential for the model’s prediction, we applied SHapley Additive exPlanations (SHAP), a method for interpreting a model’s predictions ^33^. SHAP analysis revealed that the fluorescence lifetime intervals of 2.19–2.92 ns and 5.1-5.84 ns were associated with “responders” and the intervals 2.92-3.65 ns and 3.65-4.38 ns were associated with “non-responders” (Fig. 1F). These results demonstrate that the distribution of chromatin compaction derived from measures of single nuclei fluorescence lifetime values, can predict NACT resistance in TNBC patients.

### Spatial gradients in machine learning predictions associated with transitions in chromatin compaction

Our analysis pooled all nuclei across the cores, not considering the possibility that the local organization of nuclei may contribute to the prediction of NACT response. To explore potential intra-sample spatial heterogeneity in pre-NACT cores, we extended the classification pipeline and trained a patch-based classifier (Fig. 2A, left-to-right). Each fluorescence lifetime image was partitioned into image patches, each of size ∼3 mm^2^, containing a median of 5,447 nuclei per patch (IQR: 1,595–12,276) (Fig. S3). Each patch was represented by its corresponding distribution of single nuclei fluorescence lifetimes and was assigned the NACT response for its corresponding core. We repeated the 5-Fold cross validation described above using the patches’ representations and labels. Classification performance was measured by scoring each core according to the average classification score across all of its patches, reaching a median ROC-AUC of 0.66 (maximum of 0.78) (Fig. 2B), and showing consistent interpretability with the tissue-level model (Fig. S4). To exclude the possibility that the model was biased by local nuclei density, we measured the correlation between the nuclei count in a patch and its corresponding predicted probability, which showed a negligible Pearsons’ correlation coefficient of 0.025 (Fig. S5). The degradation in classification performance from ROC-AUC of 0.735 to 0.66 was expected due to a core containing almost four orders of magnitude more nuclei than a patch. However, classification at the patch level enabled us to explore the spatial heterogeneity within a core. Mapping the patch classification scores back to space showed that most cores were characterized by spatially homogeneous classification scores (Fig. S6), potentially because core biopsies are largely derived from the tumor region with little surrounding non-tumour tissue. However, a proportion of samples exhibited spatial heterogeneity, including spatial transitions from predicted responder to non-responder regions. Exploring the fluorescence lifetime values along these spatial transitions in the same core from predicted responder to predicted non-responder showed gradual increasing fluorescence lifetime values in 3 out of the 4 spatial transitions identified (Fig. 2C-D, Fig. S7, Methods). This pattern was consistent with our SHAP interpretability of the entire core classification (compare Fig. 1F and Fig. S4). This gradual change in lifetime values did not appear in homogeneously predicted regions in the same cores and was not sensitive to the parameters used to pool the lifetime along the transition (Fig. 2C-D, Fig. S7, Methods). These results confirm our interpretation for the range of lifetime values associated with the classifier decision.

**Figure 2.**
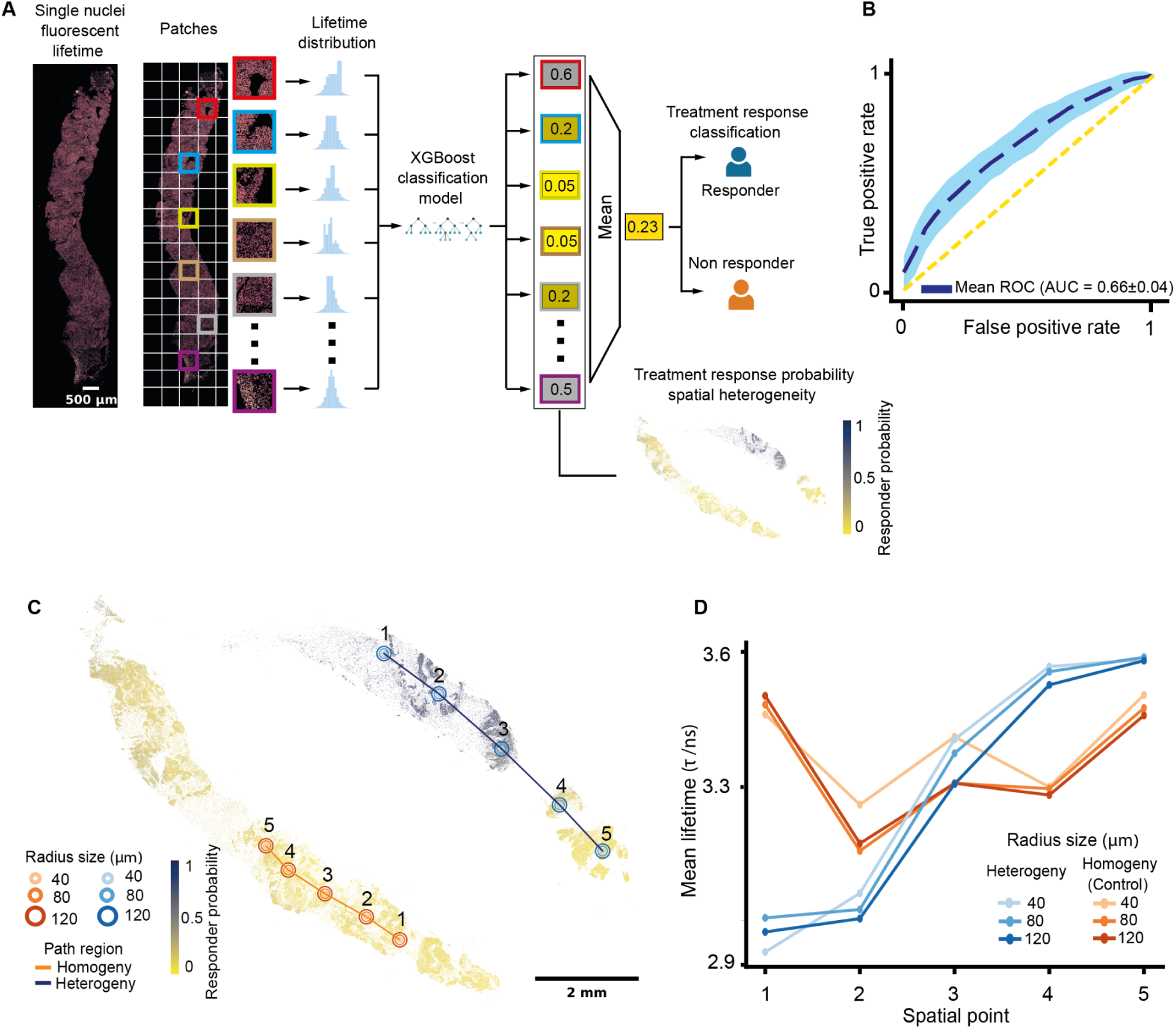
Spatial analysis of TNBC NACT treatment response prediction. **(A)** The patch-based classification pipeline. Each core image was partitioned to patches (colored boxes), each represented by its nuclei fluorescence lifetime distribution. An XGBoost classifier was trained using these patch-level distributions to classify treatment response, where each patch was assigned with its corresponding core’s label. Patch predictions were aggregated for tissue-level classification and used to explore the spatial heterogeneity in the model’s prediction. **(B)** Receiver Operating Characteristic (ROC) curve of the patch-wise XGBoost classification model. Each core was scored according to the average classification score across all of its patches. The average ROC curve (blue dashed line, ROC-AUC = 0.66) and the standard deviation (shade) across 100 different random seeds. The yellow dashed line Y = X represents random discrimination (ROC-AUC = 0.5). **(C)** Spatial visualization of the patch-level treatment response prediction (yellow = low, blue = high) for a representative tissue core. A spatial transition in the prediction from responder to non-responder is marked (blue line), with an internal control of a homogeneous non-responder prediction (orange line). These classification transition and control regions were spatially sampled with five locations (#1–5). Concentric rings around each location represented the 40 μm, 80 μm, and 120 μm radii to compute average nuclear fluorescence lifetime (circle size). **(D)** The average fluorescence lifetime values along the locations 1-5 described in panel C. Color and markers correspond to panel C.

### Spatially encoded information contributes to NACT resistance prediction

We hypothesized that incorporating information regarding the spatial organization of the nuclei would provide discriminative power and improve the prediction of NACT resistance. We firstly assessed spatial relationships in the nuclear fluorescence lifetime. For each nucleus we measured the average absolute difference in fluorescence lifetime with respect to all nuclei within a series of non-overlapping concentric 20 µm “rings” of equal widths in increasing distances with respect to the ‘central’ nucleus (Fig. 3A, Methods). Then, we calculated the correlation between the series of ring distances and the corresponding average absolute difference in fluorescence lifetime (Fig. 3B, Methods). A positive correlation coefficient indicates that the differences in lifetime values increase with the distance from the examined nucleus, implying that the lifetime values of that nucleus are more similar to the other nuclei in their close vicinity compared to other nuclei that are further away. A correlation coefficient of zero implies that there is no association between the distance from the nucleus and the difference in lifetime values, indicative of random spatial order. At the core level, we averaged the fluorescence lifetime differences across all nuclei at each distance ring and computed the correlation between these aggregated differences and the corresponding distances. The correlation coefficient showed that the vast majority of cores had correlation coefficients that exceeded 0.9 (Fig. 3C). This demonstrates that nuclei exhibit more similar fluorescence lifetime values to their neighbours compared to nuclei located further away confirming that nuclear lifetime, and hence chromatin compaction, were spatially associated.

**Figure 3.**
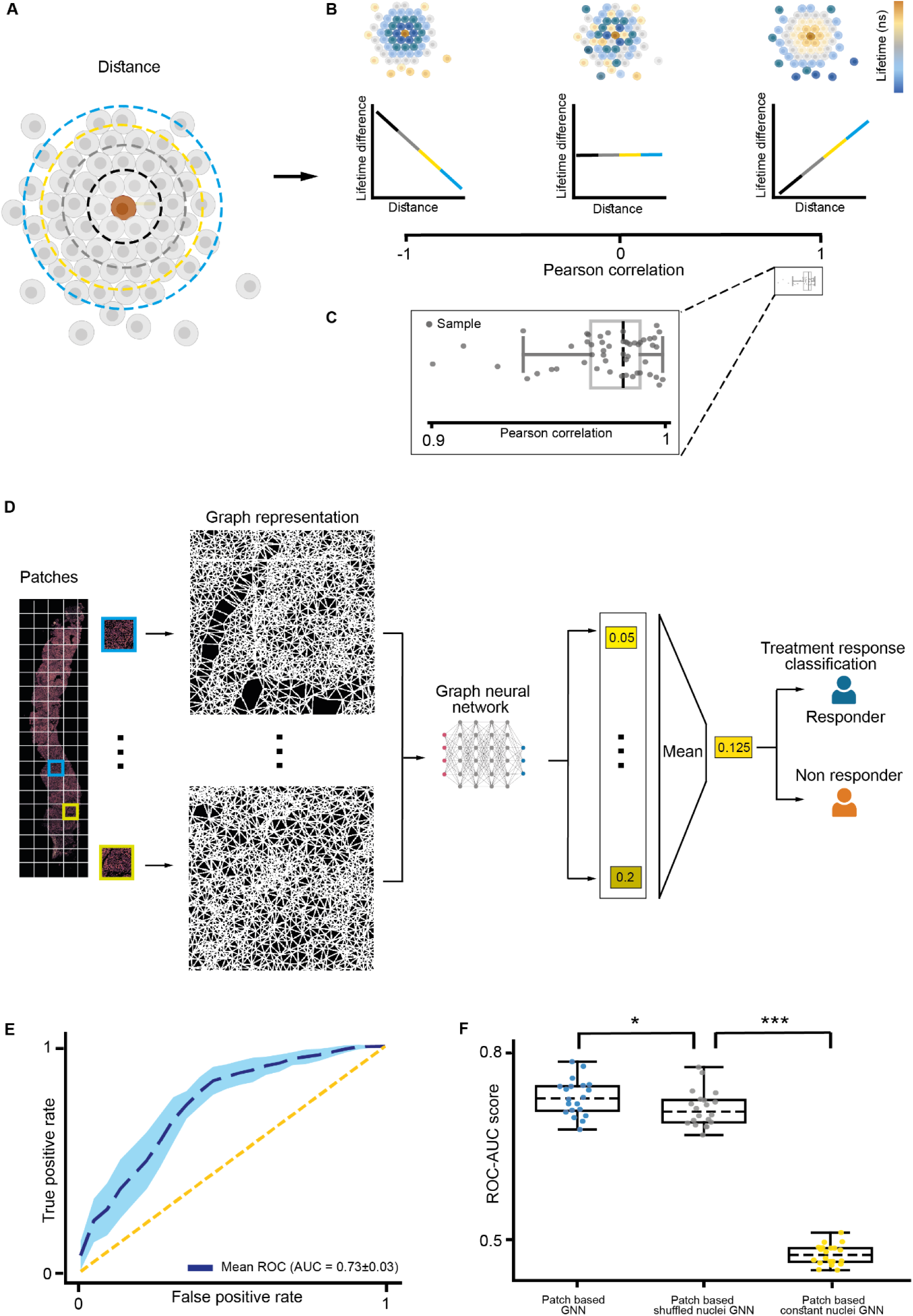
Incorporating spatial information with graph neural networks. (A-C) Spatial associations in fluorescence lifetime. (**A**) A central nucleus (brown) is surrounded by nuclei that are partitioned by non-overlapping concentric 20 µm “rigs” in increasing radia distances in respect to the central nucleus (dashed colored circles). **(B)** Illustrations of (left-to-right): negative, zero and positive correlation patterns between the rings’ distances (x-axis) and the average absolute difference in fluorescence lifetime between the central nucleus and all other nuclei in each ring (y-axis). Illustration of the spatial organization of nuclei around a central nucleus, colored according to fluorescence lifetime (top) and the corresponding plots for correlation analysis, with colored line segments (black, gray, yellow, blue) corresponding to the radii defined in panel A (bottom). **(C)** The distribution of cores (data points) with Pearson correlation coefficients between 0.9 and 1. **(D-F)** Graph Neural Network (GNN) classification. (**D**) classification pipeline. Image patches were represented as networks of single nuclei as nodes and adjacent nuclei connected by edges. A GNN was trained on these network graphs to classify treatment responses. Patch-level predictions were aggregated for core classification. **(E)** Receiver Operating Characteristic (ROC) curve of the patch-based GNN model, where each core was scored according to the average classification score across all of its patches. The average ROC curve (blue dashed line, ROC-AUC = 0.73) and the standard deviation (shade) across 20 different random seeds. The yellow dashed line (Y = X) represents random discrimination (ROC-AUC = 0.5). **(F)** Distribution of ROC-AUC across different random seeds (data points) and models (left-to-right): patch-based GNN (blue, median AUC = 0.73), patch-based GNN with shuffled lifetime values (gray, median AUC = 0.71), and patch-based GNN without lifetime values (yellow, median AUC = 0.48). One-sided Mann–Whitney U test with Benjamini-Hochberg FDR correction for multiple comparisons, where * = adjusted p-value < 0.05 and *** = adjusted p-value < 0.001.

Our machine learning models (Figs. 1-2), did not consider the spatial organization of the nuclei in the tissue. To encode this spatial information into a predictive model, we represented image patches as networks, or graphs, of adjacent nuclei and trained a graph neural network (GNN) model to implicitly incorporate the nuclei fluorescence lifetime along with these spatial dependencies between adjacent nuclei. The graph was represented with nuclei as nodes, their fluorescence lifetime values as node features, and with spatial adjacency of immediate neighboring nuclei as edges. GNNs iteratively aggregate node features across edges through a process called “message passing” ^34^. In message passing, each node (nucleus) combines features from nodes connected to it by an edge to update its own feature toward minimizing a target function (“loss”), which in our case is the treatment response prediction. We used 13,893 image patches of size ∼3 mm^2^ each, pooled from the 53 cores to train a GNN to predict the treatment response. For each core, the predictions at the patch level were accumulated and the average prediction was reported. The full classification pipeline is depicted in Fig. 3D and in the Methods. To ensure robustness and account for variability, we trained 20 different models with different random seeds. The corresponding 20 GNN models reached a median ROC-AUC of 0.73 (maximum of 0.79) (Fig. 3E). GNNs trained on the same networks after randomly shuffling the fluorescence lifetime values among nuclei within each patch, reached a mildly reduced performance indicative of the contribution of spatial lifetime information to the predictive performance (Fig. 3F, Methods). The performance of GNNs trained on the network structures without lifetime values dropped to the level of random prediction indicating that nuclear position is not predictive without encoded fluorescence lifetime (Fig. 3F). These results establish that the spatial arrangement of nuclei chromatin compaction is predictive of NACT resistance.

### Chromatin compaction is preserved in tissue state transitions from pre- to post-NACT

We hypothesized that the nuclear fluorescence lifetimes in the same TNBC patient tissues pre (core) and post (resection) treatment were associated, maintaining a chromatin compaction “signature” throughout the disease progression (Fig. 4A). Direct spatial comparison between the core and the resection samples was not possible because they were taken several months apart, and because the resection specimens were substantially larger and more heterogeneous than the core biopsies. Instead, we devised an indirect approach to measure similarities between matching core-resection pairs. First, for each core and resection, we recorded the median fluorescence lifetime across all nuclei. Evaluating the distribution of these median fluorescence lifetimes revealed that responder and non-responder cores were not significantly different; however resection samples had significantly higher median lifetime values than responder cores (Fig. 4B). To more directly investigate the intra-patient progression, we applied a matched pair ranking process ^35^ to ask whether the fluorescence lifetime patterns are preserved between pre-treatment cores and their corresponding post-treatment resections.

**Figure 4:**
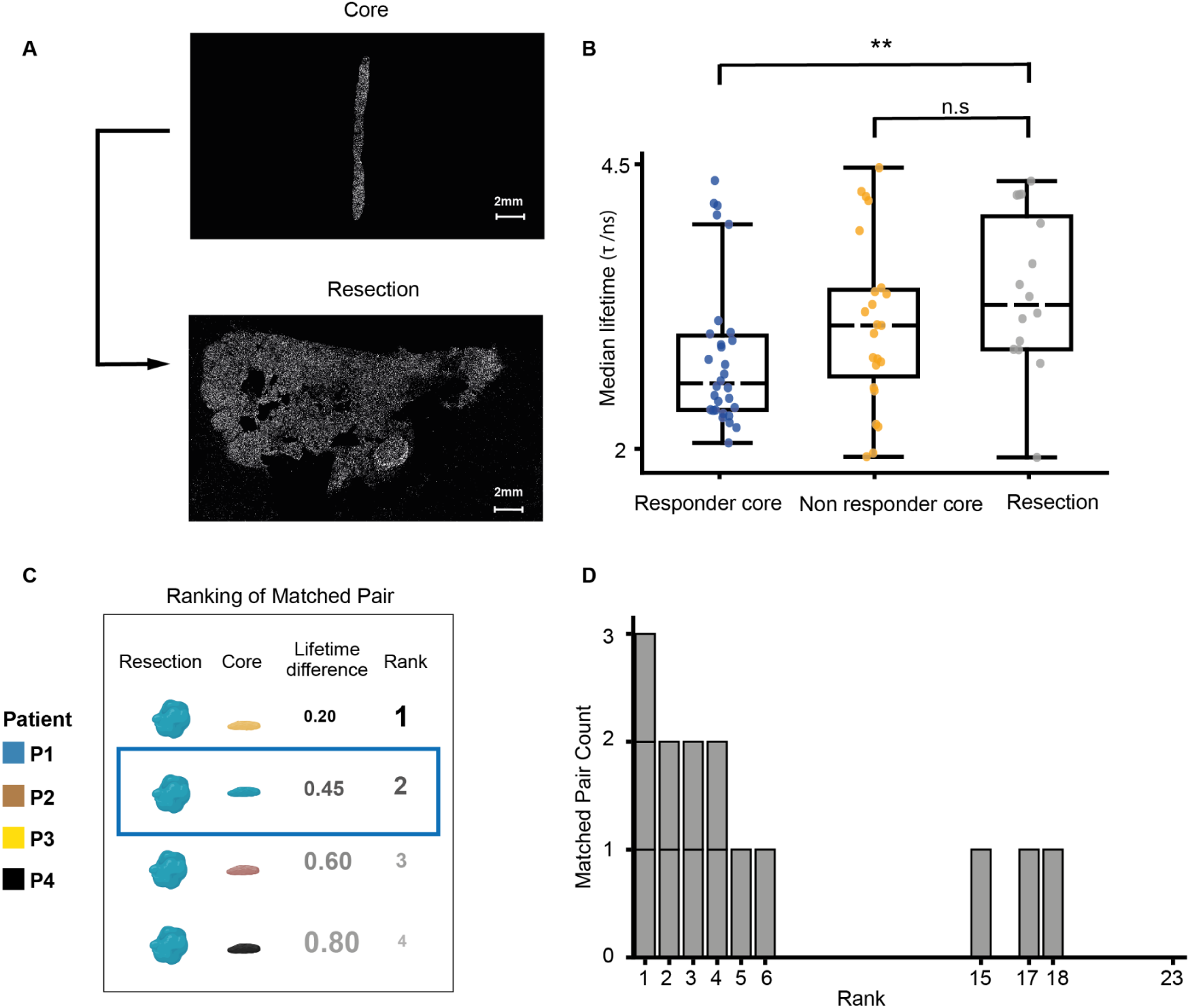
Fluorescence lifetime association between cores and matched resections. **(A)** Representative fluorescence lifetime images of a pre-treatment core biopsy (top) and corresponding post-treatment surgical resection (bottom) from the same non-responder patient. Scale bar = 2 mm. **(B)** The distribution of median fluorescence lifetime values for core biopsies from responder patients (blue), core biopsies from non-responder patients (orange), and resections from non-responder patients (gray). Each data point represents a single patient’s core or resection. The black dashed line indicates the group median. Two-sided independent t-tests with Benjamini-Hochberg FDR correction for multiple comparisons, were used to test the null hypothesis (H₀: equal median fluorescence lifetimes between groups) attaining adjusted p-value = 0.006 (**) for responder cores versus non-responder resections, and non-significant adjusted p-value = 0.147 (n.s) for non-responder cores versus resections. **(C)** Depiction of an example of the matched pair ranking process of the resection and its corresponding non-responder core. In this example, colors denote patients (P1–P4) and the blue box highlights the matched core-resection pair. The resection was compared to its matched core from the same patient and to all non responder cores from other patients. The similarity between the resection and its matched core was ranked relative to the similarity of that resection to all other non responder cores where a rank of 1 indicates the highest similarity. **(D)** Distribution of ranks for 14 matched core-resection pairs using a similarity score based on the absolute difference in their median fluorescence lifetime. The x-axis shows the rank of the matched core-resection pair (1 = most similar), and the y-axis indicates the number of matched core-resection pairs that received each rank.

This was approached by devising a measurement for how well each resection was matched to its corresponding non-responder core (Fig. 4C). Firstly, for each matched core-resection pair originating from the same patient, we calculated a similarity score based on the absolute difference between their median fluorescence lifetime values. Secondly, for each resection, we calculated the similarity to each one of the core samples from the other non responder cores. Finally, we ranked the similarity between the resection and its matched core in respect to the resection’s similarity to the other non responder cores. The rank ranged from 1 to 𝑁 = 23, where 𝑁 represents the total number of non responder cores compared for that resection. A rank of 1 indicates that the matched core from the same patient was the most similar to its corresponding resection, while a rank of 23 indicates it was the least similar among all cores. 10/14 (∼71%) of the matched core-resection pairs were ranked within the top 5 matches against non-responder cores (Fig. 4D), three folds above the expected random matching probability of 5/23 (∼22%) (Methods). These results indicate that the chromatin compaction signature is preserved in the tissue state transition from core to resection.

## Discussion

Triple-negative breast cancer (TNBC) remains a major clinical challenge due to its molecular heterogeneity, poor prognosis, and limited therapeutic options. Here, we introduce chromatin compaction - quantified by fluorescence lifetime imaging microscopy (FLIM) - as a previously underexplored spatial predictor of therapeutic response. By reducing the complexity of nuclear organization into a single quantitative lifetime metric, we show that tumors resistant to neoadjuvant chemotherapy (NACT) exhibit more open chromatin states that are spatially coordinated within the tissue, and which persist during NACT treatment.

These findings establish chromatin organisation as a quantifiable spatial discriminator of therapeutic resistance.

Previous studies have shown the chromatin state can be used to distinguish between different histological annotations ^17,18^, cancer cell states ^17^, disease stages ^18^ and breast cancer subtypes ^19^ . We report here that chromatin structure can also predict therapeutic response, demonstrating that the present chromatin state contains predictive information about future drug resistance. Notably, we represent a nuclear chromatin state with a single value, in contrast to other studies that used higher resolution and/or alternative measurements that describe nuclear organisation ^17,18,21,36,37^. Our representation of per nuclei fluorescence lifetime was sufficient to predict resistance to chemotherapy, to uncover intra- and inter-sample spatial heterogeneity, reveal that the spatial arrangement of nuclei contributing to response, and to demonstrate the preservation of chromatin signatures between pre- and post-treatment biopsies in resistant patients.

Plasticity in chromatin organisation has been previously linked to resistance to chemotherapy by transient chromatin compaction following treatment, followed by decondensation during recovery ^21^ , though the specific role of chromatin compaction in chemoresistance remains contested. Some studies suggest that condensed chromatin confers protection by limiting DNA damage ^22^ and repressing DNA repair genes ^23^. Other studies propose that more open chromatin facilitates resistance through transcriptional flexibility ^21^ and alters sensitivity to targeted agents ^24,25^ . Our results propose that using chromatin state as a bulk readout across the tissue does not predict chemotherapy resistance. Rather, the heterogeneity and spatial association of cells with less compact chromatin can predict response. The adaptation of open chromatin state in some cell types may reflect disruption of specific nanodomains, thereby promoting transcription of important drug-resistance genes. This is likely due to extensive changes in post translational modification of histones that weakens internucleosomal associations and ’relaxes’ chromatin organisation. Indeed, therapeutic agents that reduce histone modifications have been proposed as solutions for drug resistant TNBC ^38^. Further mechanistic interpretation of the spatial topography of the open chromatin states we have uncovered into TNBC NACT resistance would be aided by corresponding multimodal data ^39^ encoding cell type specific markers and extracellular components that may contribute to the features we report. Nevertheless, results emphasize the physiological signal encapsulated in chromatin fluorescence lifetime representation could be extended to other cancer types and integrated to enrich interpretation of orthogonal multiplexed spatial omics techniques.

## Acknowledgements

This research was funded by the Wellcome Leap ΔTissue program. Reut Mealem and Leor Ariel Rose were supported by the Kreitman STEM scholarship. We thank Orit Kliper Gross for critically reviewing our GitHub repository. We thank Dan Levy and Noga Levy for critically reading our manuscript. We thank Anisha Kubasik-Thayil, Nicholas Anthony and Andreas Boden of the Microscopy Innovation Centre at King’s College London and Rocco D’Antuono of the Crick Advanced Light Microscopy facility at the Francis Crick Institute for assistance with FLIM data acquisition. We also thank the King’s Health Partners Cancer Biobank for assistance with tissue sectioning and clinical metadata retrieval.

## Author Contribution Statement

RM, TP and LAR contributed equally to this work. MP and AZ conceived the study and acquired funding. RM and LAR developed the computational methods, and analyzed the data. TP optimised sample preparation workflow, prepared samples for imaging and acquired the data. TP and SM preprocessed the data. All authors interpreted the data. RM, LAR, MP and AZ drafted the manuscript. All authors edited the manuscript and approved its content.

AZ mentored RM and LAR. MP mentored TP and SM.

## Methods

### Patient cohort and preparation of tissue sections

Tissues were collected as 1 mm core needle biopsies from patients pre-NACT or surgical resections post-NACT (more detailed cohort information can be found in ^40^). Our cohort consisted of cores for 30 responders and for 23 non-responders, and resections for 14 of the non-responders. Formalin fixed paraffin embedded (FFPE) TNBC tissue sections were prepared for staining and imaging by incubating slides on their side in a 65°C oven for at least 2 hours to melt paraffin wax. Following this, a stepwise rehydration of slides was accomplished by submersion of slides for 2x10 minutes per cycle in xylene, 100% ethanol, 70% ethanol (v/v with distilled water), 50% ethanol and distilled water. Antigen retrieval was achieved using pH 6 citrate buffer (24.88 g sodium citrate and 21 g citric acid monohydrate in distilled water to total volume of 2 L) at 95°C for 30 minutes. Following antigen retrieval, samples were treated with 1mg/mL sodium borohydride in tris-buffered saline (TBS) for 10 minutes to reduce autofluorescence and washed thrice for 3 minutes in TBS on a rocker.

Samples were then permeabilised by treating with 0.2% tween in TBS (TBST) for 30 minutes, washed thrice for 3 minutes and then blocked using 5% (w/v) bovine serum albumin (BSA) in TBST for 30 minutes before addition of 1:500 SiR-DNA for 2 hours in a dark, humidified chamber before washing three times in TBS for 10 minutes and mounting a coverslip on top of the section using Fluorosafe.

### Validation of SiR-DNA probe in cultured cells

HeLa cells (ATCC) were plated onto glass coverslips in 12-well tissue culture plates at a density of 50000 cells/mL and cultured in EMEM supplemented with 10% FBS, 1% penicillin-streptomycin and 1% L-glutamine for 24 hours. Cells were cultured in the presence of 1:500 SiR-DNA (Spirochrome) and 1:1000 verapamil (Spirochrome) for 24 hours in 0%, 2%, 10% and 20% dextrose solutions (w/v) in EMEM to induce osmotic shock and chromatin compaction ^30^. After 24 hours culture, samples were washed thrice in PBS and fixed in 4% PFA for 10 mins, followed by three more PBS washes and mounted onto SuperFrost Plus microscope slides using Fluorosafe mounting media.

### Fluorescence lifetime imaging microscopy (FLIM)

FLIM was carried out using a Leica Stellaris FALCON microscope (Leica Microsystems) equipped with a white light laser (WLL). Samples were imaged using a 10x air objective (0.4NA) with additional 2x magnification. SiR-DNA was excited using 11.98% laser power at 651 nm and the emitted signal was gated at 658-760nm and captured on a HyDS2 detector. LAS Navigator was used to define the boundary of the tissue sections and build a tiled series of individual images to cover the entire section area. Tiled regions were then stitched into single images using LAS X software. To permit accumulation of sufficient photons for fitting of decay curves, a frame repeat of 5 was applied to each tile acquired and a minimum average signal of 1000 counts was required for each tile to allow for robust lifetime fitting. For sucrose-treated HeLa cell validation imaging, a minimum of five fields of view were imaged per condition per repeat, with three independent repeats imaged in total. Following imaging, data were fitted in LASX using an instrument response function (IRF) for a mono-exponential decay curve model to estimate the fluorescence lifetime τ (the excited state decay time) for each pixel, reported in picoseconds. Data was exported for downstream analysis using the LAS X software to output merged TIFFs with a spatial resolution of 1.139 μm per pixel. Each TIFF contains the fluorescence intensity image, which indicates total photon counts per pixel and lifetime-encoded image.

### FLIM preprocessing workflow

To ensure consistent intensity range across cores, the fluorescence intensity images were percentile-normalized using CSBDeep’s ^41^ normalize function with default settings percentiles. Segmentation was then performed on these normalized intensity maps with the StarDist2D model pretrained on the “2D_versatile_fluo” dataset ^31^, with defaults parameters except for scale = 3 and for non-maximum suppression threshold = 0.6 to reduce overlapping detections. We visually inspected segmentation outputs in Napari ^42^ and applied size filters to remove objects below the 5th percentile or above 100 pixels (≈114 µm²), excluding under 10% of nuclei (Fig. 1C). This approach ensured that only accurately segmented, reliably measured nuclei were considered for the downstream fluorescence-lifetime analyses. Using each segmentation mask together with its corresponding fluorescence lifetime channel, we extracted per-nucleus readouts and compiled these into a single-nuclei table for downstream analysis. Readouts included the nucleus unique label, centroid coordinates, fluorescence lifetime, shape features (eccentricity and size) and local density. Nuclei with fluorescence lifetime above 13 nanoseconds were filtered out as they represented rare outliers and because the imaging system’s calibrated dynamic range extends to 13 ns.

### Machine learning NACT-resistance prediction using tissue-wise distribution of single nuclei chromatin compaction

Each fluorescence lifetime image was converted into a distribution of single nuclei fluorescence lifetime by dividing the 0–13 nanoseconds range into equal-width bins and computing the fraction of nuclei falling into each bin. We evaluated five binning schemes (10, 18, 23, 34 and 42 bins) to identify the optimal resolution. These full distributions served as feature inputs for an XGBoost ^32^ classifier tasked with discriminating NACT response of tissue cores. Due to the limited number of patients in the cohort, we trained XGBoost classification models to predict response with a leave-one-core-out cross-validation strategy using scikit-learn’s ^43^ LeaveOneGroupOut. In each fold one core was designated as “test” and all other cores were used for training. To ensure robustness and account for stochastic variability, we repeated model training (and then testing) across 100 different random seeds, providing a comprehensive assessment of performance stability. Hyperparameter tuning for each model was conducted within each training fold using the Optuna ^44^ framework (50 trials per fold), optimizing for ROC AUC. The search space for XGBoost included maximum tree depth (3–10), learning rate (0.01–0.3, log scale), subsample ratio (0.5–1.0), column subsampling (0.5–1.0), γ (0–10), number of estimators (50–500), L1 and L2 regularization terms (1–10, log scale), and scale_pos_weight (1–10). To confirm that these results were not due to overfitting, we shuffled the NACT response labels, retrained the classification models (using the same fixed hyperparameters from our median-seed classification model trained on the true labels) and recorded the shuffled ROC-AUC. Repeating this process 1000 times and assessing how many different shufflings induced performance equal or better than the median ROC-AUC observed with the true (i.e., non-shuffled) labels. The empirical p-value was calculated as the proportion of permuted runs with an AUC greater than or equal to that obtained under the true labels. Among the five binning resolutions evaluated, the 18-bin configuration achieved the highest classification performance (Fig. S8) and was selected for downstream analysis. To pinpoint which interval ranges of fluorescence lifetime values were the most influential for the model’s prediction, we applied SHapley Additive exPlanations (SHAP), a method for interpreting a model’s predictions ^33^.

### Machine learning NACT-resistance prediction using patch-based distribution of single nuclei chromatin compaction

Patch-wise representations were generated similarly to the tissue-level distribution of single nuclei chromatin compaction, but at a patch scale. Each fluorescence lifetime image was divided into overlapping square image patches of 1500×1500 pixels (≈1708² μm²), with 75 % overlap between adjacent patches (Fig. 2A). Patches containing fewer than 50 nuclei were discarded. For each remaining patch, we computed the fluorescence-lifetime distribution using 18 equal-width bins over 0–13 nanoseconds and expressed each bin’s count as a fraction of the patch’s total nuclei. Because ground-truth labels exist only at the tissue (core) level, each patch inherits its parent core’s NACT response label. Model training and evaluation reused the XGBoost pipeline from the tissue-level analysis (100 random seeds per fold, Optuna hyperparameter tuning and median-seed permutation testing). To leverage the larger number of patches and accelerate training, we replaced leave-one-core-out with grouped stratified 5-fold cross-validation utilising scikit-learn ^43^, where 20% of the patches were used as the test, and the remaining 80% were used for training. We ensured that patches from the same patient were kept in the same fold, preventing data leakage. To obtain tissue level prediction, patch-level predictions were then aggregated by averaging. The entire pipeline was repeated using a smaller and a larger patch size of 1000×1000 and 2000×2000 pixels (≈1139² and 2278² μm² correspondingly) to validate patch size sensitivity. The sensitivity analysis showed no significant difference in the model’s performance (Fig. S9).

### Analysis of spatial gradients in the classification prediction

To assign our patch-based predicted treatment response probabilities at single-nuclei resolution, we averaged the predicted classification probability of all image patches that overlapped with its spatial location. We manually selected continuous regions in cores that showed gradual transitions from patches predicted as responder to patches predicted as non-responders (Fig. S7). We identified four cores with such transitions in our cohort. As internal control we selected a continuous region in the same core that was homogeneously predicted as non-responder. Each of the classification transitions and the control region were manually spatially sampled with five locations that were approximately equally spaced using Napari ^42^. To measure changes in the fluorescence lifetime during these transitions and their corresponding controls, we averaged the lifetimes of all nuclei within concentric circular regions centered at each of the manually selected locations around the transition/control with radii of 40, 80, 120 μm.

### Measuring spatial relationships in nuclear fluorescence lifetime

For each nucleus, we generated six non-overlapping concentric 20 µm rings of equal widths in increasing distances with respect to the ‘central’ nucleus (Fig. 3A). In each ring, we computed the ‘central’ nucleus average absolute difference in fluorescence lifetime with respect to all nuclei within the ring. For each core, we pooled the per-nucleus average difference in fluorescence lifetimes across all nuclei yielding a core-wise average absolute difference in fluorescence lifetime for each ring. This analysis resulted in 6 values encoding the average absolute change in the lifetime as a function of distance across all nuclei in a given core. The correlation between these average absolute differences in lifetime and their corresponding distances was used as a measure for spatial relationships in the nuclear fluorescence lifetime. A positive correlation coefficient indicates that the differences in lifetime values increase with the distance from the examined nucleus, implying that the lifetime values of that nucleus are more similar to the other nuclei in their close vicinity compared to other nuclei that are further away. A correlation coefficient of zero implies that there is no association between the distance from the nucleus and the difference in lifetime values, indicative of random spatial order, and a negative correlation implies that lifetime values of nuclei in a core are more similar to nuclei that are further away than to nuclei in their close vicinity (Fig. 3B).

### Graph neural network NACT-resistence prediction

To incorporate spatial context to our predictive models we defined graphs where individual nuclei served as nodes and the fluorescence lifetime value of each nucleus was used as the node feature. Edges in the graph were defined by the spatial adjacency of immediate neighboring nuclei using delaunay triangulation, implemented via scipy.spatial.Delaunay, which approximates a voronoi diagram over the spatial coordinates of segmented nuclei centroids. To ensure biological relevance we excluded edges exceeding euclidean distance of ∼34 μm (30 pixels). Limited sample size (53 cores) and ∼130,000 nuclei per core made it not feasible to construct graphs from the entire core due to the “curse of dimensionality”.

Therefore, we constructed graphs at the image patch-level, thus providing sufficient instances for training to reduce overfitting. Image patches were generated by dividing each fluorescence lifetime image into overlapping square image patches of 1500×1500 pixels (≈1708² μm²), with 75 % overlap between adjacent patches, followed by exclusion of patches containing fewer than 50 nuclei (Fig. 3D). The graph representation of each patch was converted to a torch_geometric.data.Data object to enable training with PyTorch Geometric (PyG) and each patch-level graph was assigned with its core’s NACT response label. For graph neural network training, we applied grouped stratified 5-fold cross-validation with scikit-learn, where 20% of the patches were used as the test, and the remaining 80% were used for training. In addition, 10% out of the training patches were used for validation during training. To prevent data leakage, all patches originating from the same patient were exclusively assigned to either the training, validation, or test sets. To assess robustness to stochastic variability we repeated model training (and then testing) across 20 random seeds. A graph attention network (GAT) ^45^ was employed for graph-based patch classification, due to its ability to incorporate both node features and graph topology using attention-based weighting of neighboring nodes. The model consisted of three hidden layers with sizes 64, 128, and 64, respectively, each with a single attention head, generating node embeddings.

These embeddings were then aggregated into a graph-level representation using max pooling, followed by two fully connected layers with sizes 64 and 1 yielding a single output node for binary classification. Training was performed over 50 epochs using the AdamW optimizer, with L2 regularization (weight decay) set to 0.0001 and learning rate equal to 0.0001. In addition hyperparameter tuning was performed across several configurations, where most parameters were kept constant, and a subset was varied to evaluate performance. The dropout rate was tested at 0.1 and 0.05, batch size at 16 and 32 (Fig. S10). Based on model performance, we selected for a batch size of 32, dropout rate of 0.1. As a sensitivity analysis we repeated the entire pipeline using a smaller and a larger patch size of 1000×1000 and 2000×2000 pixels (1708² and 2278² μm² correspondingly) showing no significant difference in the model’s performance (Fig. S10). To examine the contribution of the spatial information, we trained two additional models as control. In the first control, we randomly shuffled the fluorescence lifetime values among nuclei within each patch, disrupting the spatial arrangement while preserving the global distribution of lifetimes. In the second control, we used the same graph structures, assigning a constant lifetime value of 0 to all nuclei, preserving spatial organization but removing lifetime information.

### Matched pair ranking

To statistically assess the association between resections and their corresponding non-responder cores we adapted ^35^ to devise a matched pair ranking process. Specifically, each core/resection was represented by its median fluorescence lifetime across all nuclei. For each resection we calculated the absolute difference between the resection and each of the non-responder cores’ median fluorescence lifetime. Next, we ranked the absolute difference between each resection and its matched core in respect to all other 22 non responder cores.

The rank for each core ranged between 1-23. A rank of 1 indicates that the matched core from the same patient was the most similar to its corresponding resection, while a rank of 23 indicates it was the least similar among all cores.

## Data Availability

Raw FLIM images and metadata are publicly available from the BioImage Archive ^46,47^ at https://doi.org/10.6019/S-BIAD2418 under accession number S-BIAD-2418. Metadata includes sample type, sample preservation, clinical outcome, pre/post-NACT matching and RCB score. Intermediate data products can be reproduced from the raw inputs using the source code provided in our repository. All datasets required to recreate the figures presented in this article are included as supplementary files and can be recreated using the source code.

## Code availability

Source code is publicly available at github.com/zaritskylab/TNBC-SPATIAL-CHROMATIN-COMPACTION.

---

**Figure S1.**
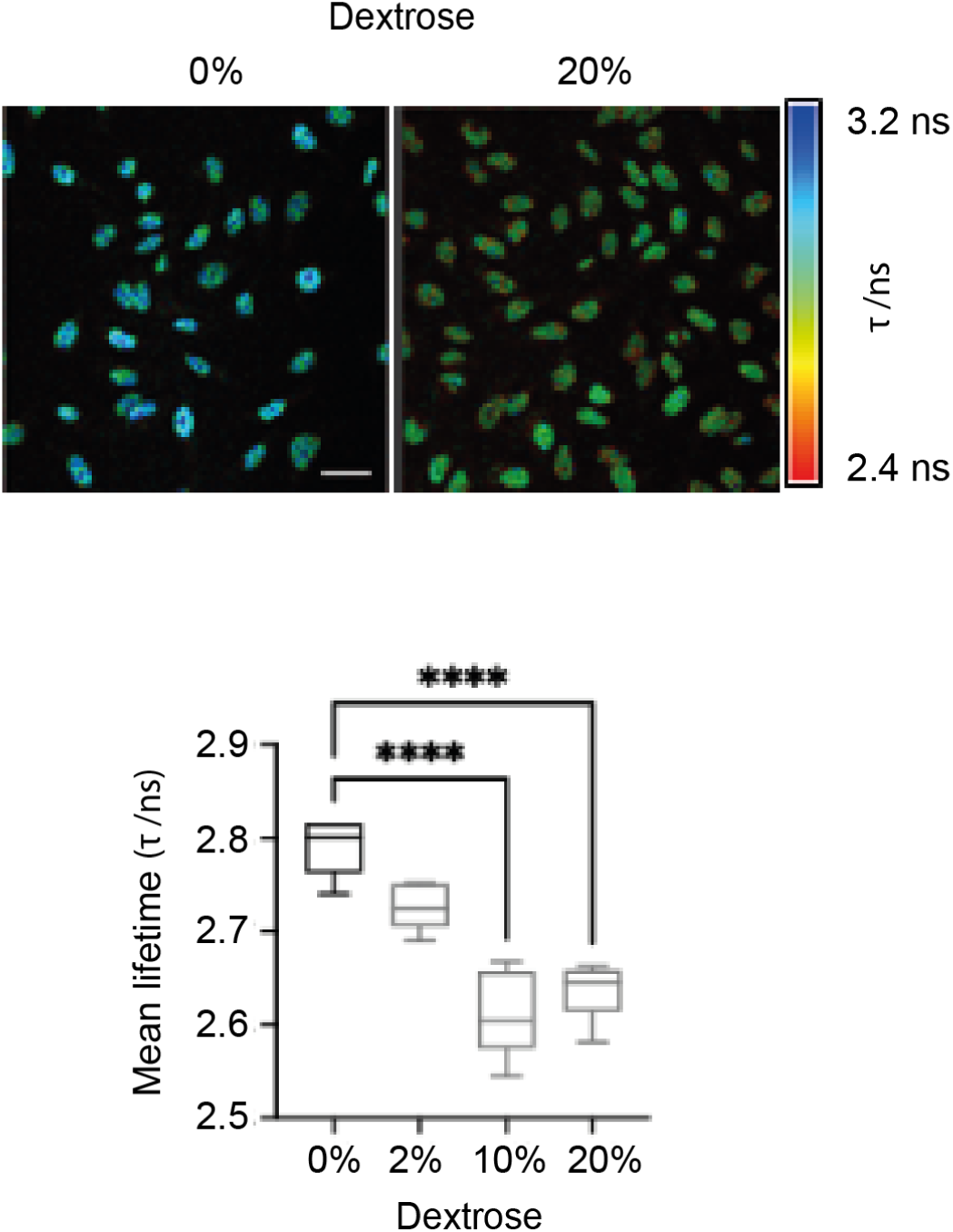
**Validation of SiR-DNA as a reporter for chromatin compaction**. Representative images of HeLa cells stained with SiR-DNA and treated with normal growth media (o%) or 20% dextrose for 24h followed by fixation and FLIM imaging. Scale bar 10 mm. Graph shows quantification of fluorescence lifetime from cells treated with increasing dose of dextrose. N=5, averages shown as line with 95th percentile. One-way ANOVA with Tukey’s HSD test; **** = p < 0.0001.

**Figure S2.**
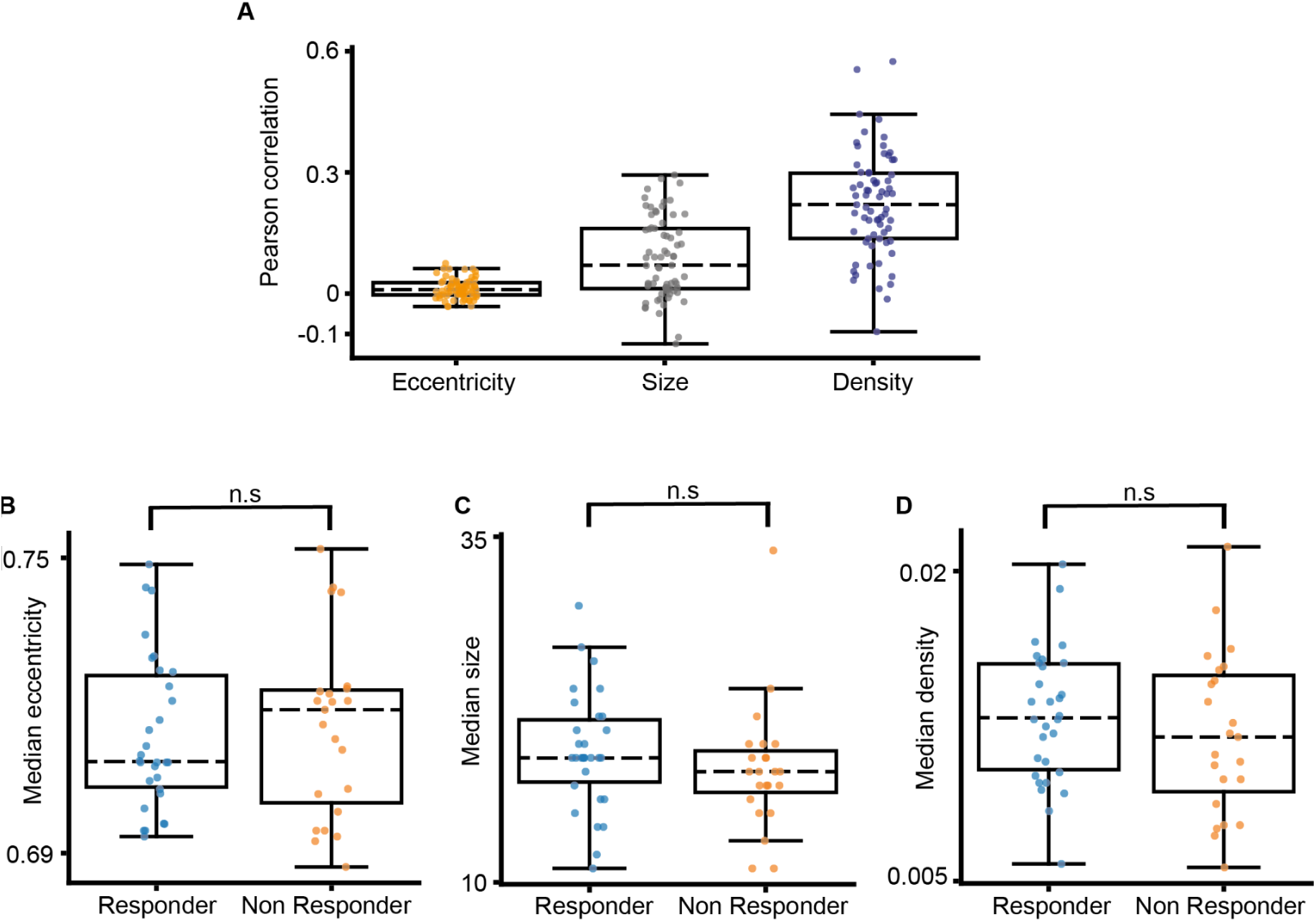
Analysis of nuclei shape and local density. Throughout this figure, each data point represents a measurement or correlation across all nuclei within a single core. (**A**) The distribution of the patient-level Pearson correlations between the fluorescence lifetime and each shape/local density feature across all nuclei for each core (left-to-right): nuclei eccentricity (orange, median correlation = 0.009), nuclei size (gray, median correlation = 0.07), nuclei local density (blue, median correlation = 0.22). (**B-D**) The distribution of the median nuclei eccentricity (B), size (C) and local density (D) across all nuclei of responder versus non-responder cores. Two-sided independent t-tests (α = 0.05) did not reject the null hypotheses (H₀: responder median = non-responder median). (B) Eccentricity, p = 0.723. (C) Size, p = 0.302. (D) Local density within a 40 µm radius, p = 0.383.

**Figure S3.**
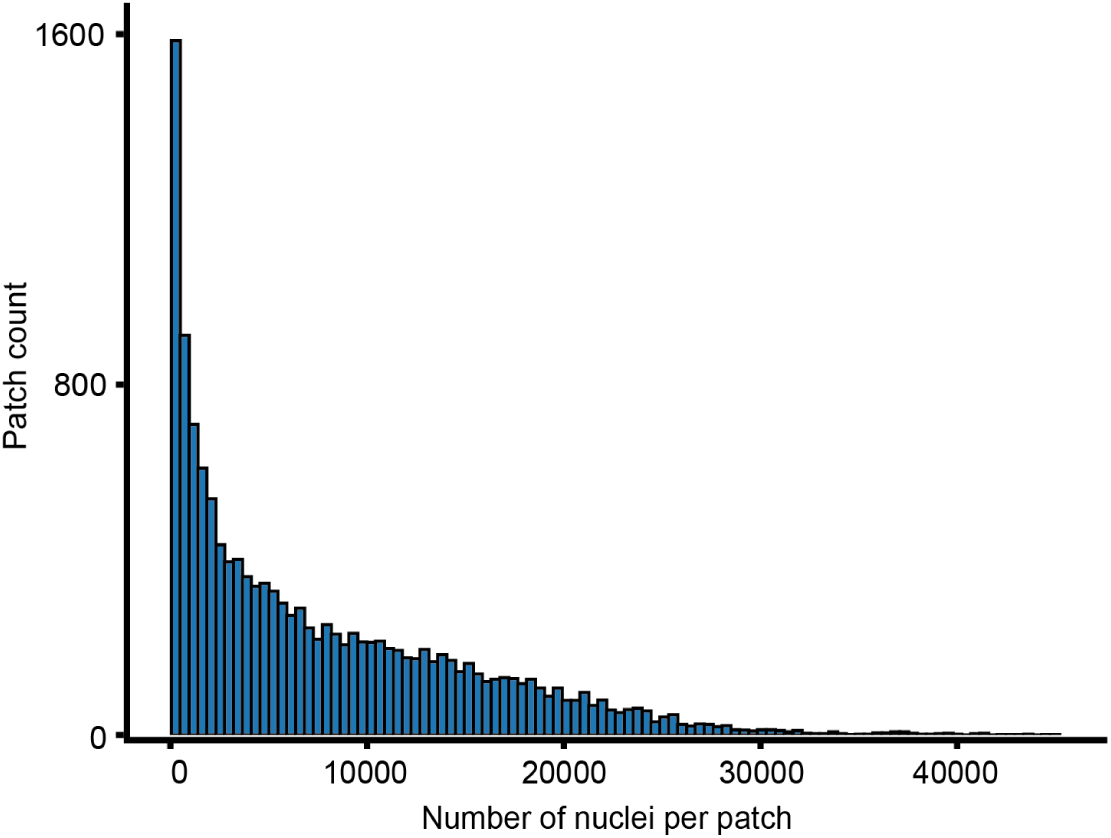
Histogram of the number of nuclei per patch. The histogram shows the number of image patches (y-axis) for each value of nuclei count per patch (x-axis).

**Figure S4.**
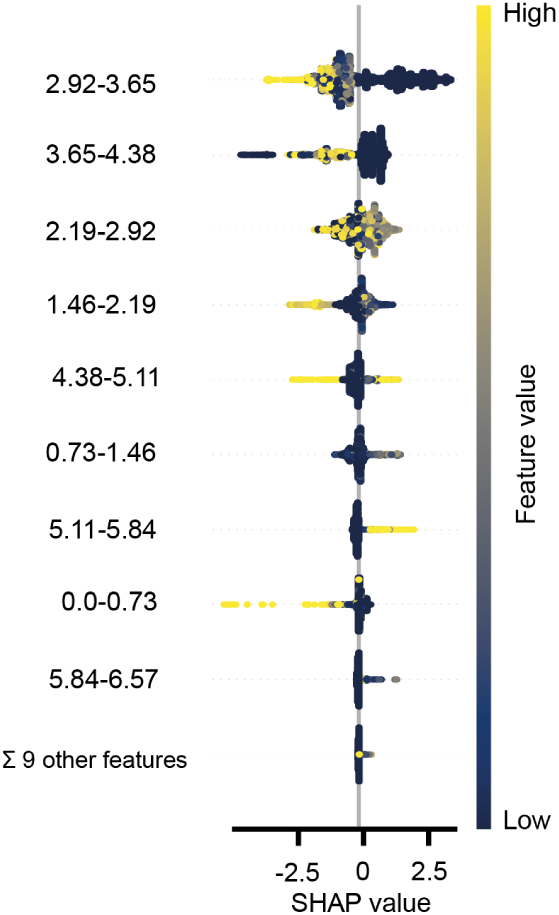
**SHapley Additive exPlanations (SHAP) analysis for the top-ranked features in the XGBoost fluorescence lifetime patch-based model**. Each horizontal feature (e.g., 2.92-3.65) represents a bin in the fluorescence lifetime distribution, with the range indicating the lifetime interval (in nanoseconds). Positive SHAP values indicate features contributing to responder predictions, while negative SHAP values correspond to features contributing to non-responder predictions. Each dot represents a patient’s SHAP value while its coloring reflects feature values (the normalized frequency of nuclei in each bin), with yellowish representing high feature value and bluish representing low feature value.

**Figure S5.**
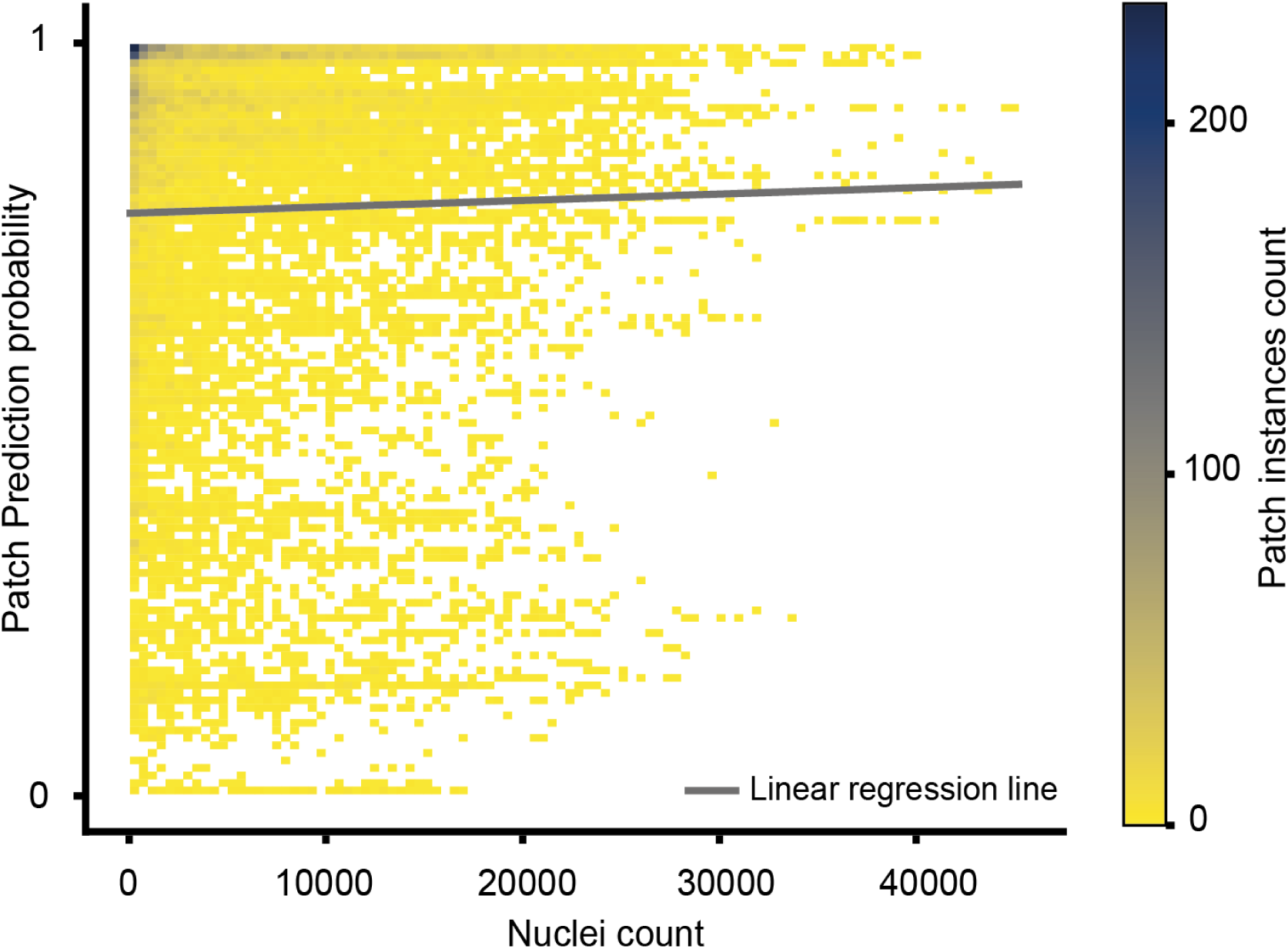
Heatmap of nuclei count versus patch prediction probability. 2D histogram where each bin represents the nuclei count in a ∼3mm^2^ patch (x-axis) and the patch-based model corresponding prediction probability (y-axis). Color represents the patch instance counts in each histogram bin, with warmer colors corresponding to lower values. The gray line represents the linear regression fit between nuclei count and patch prediction probability with a Pearson correlation coefficient of 0.025 (p = 0.0032).

**Figure S6.**
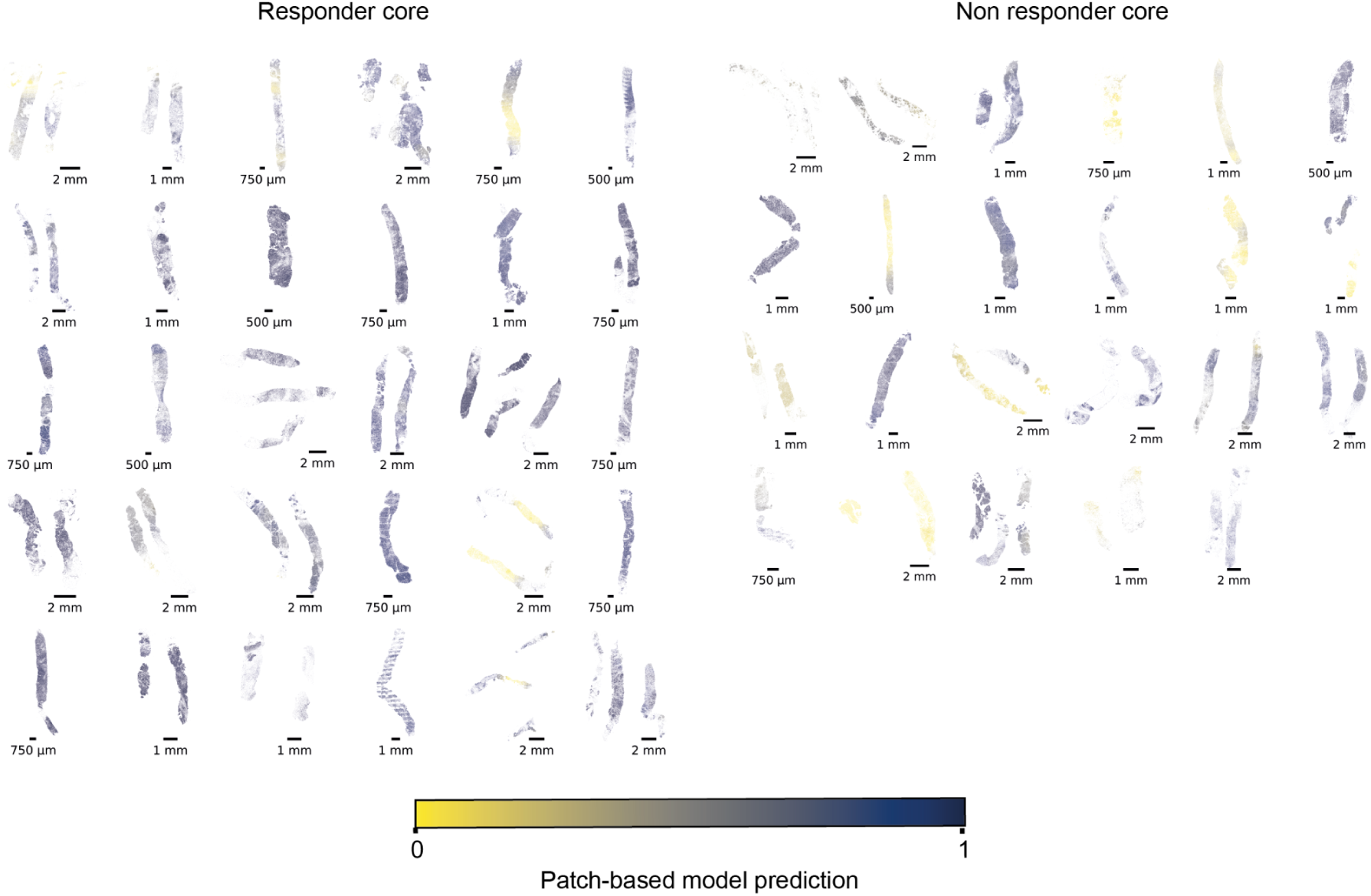
Visualization of the patch-level treatment response prediction for the entire cohort. Each panel represents a tissue core, with colors indicating patch-based treatment response predictions (yellow = low, blue = high). Left: responder cores; right: non-responder cores.

**Figure S7.**
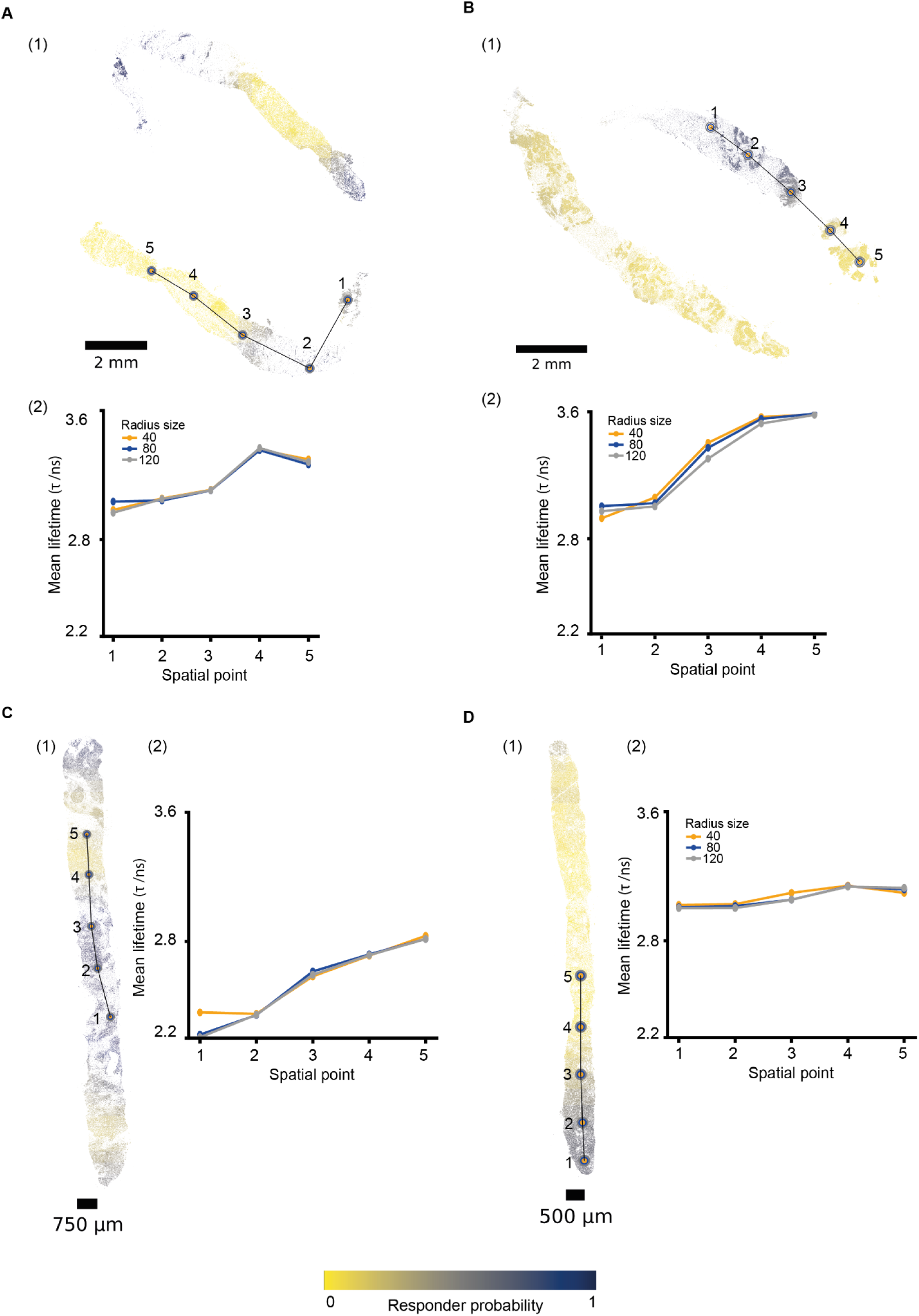
Spatial transitions in the patch-based model predictions. (A–D) Four representative tissue samples from different patients. (1) Spatial transition trajectory from responder (blue) to non-responder (yellow) predictions based on the patch-level treatment response probabilities (yellow = low, blue = high) . Along each trajectory, five spatial points were selected to represent the continuous transition. Colored circles in orange, blue, and gray around each point represent radius sizes of 40 μm, 80 μm, and 120 μm, respectively, used to compute the average nuclear fluorescence lifetime. (2) Average fluorescence lifetime values along the trajectories shown in (1), computed for three neighborhood radii (40, 80, 120 μm). Each spatial point represents the average of nuclei-level fluorescence lifetime values within the specified radius.

**Figure S8.**
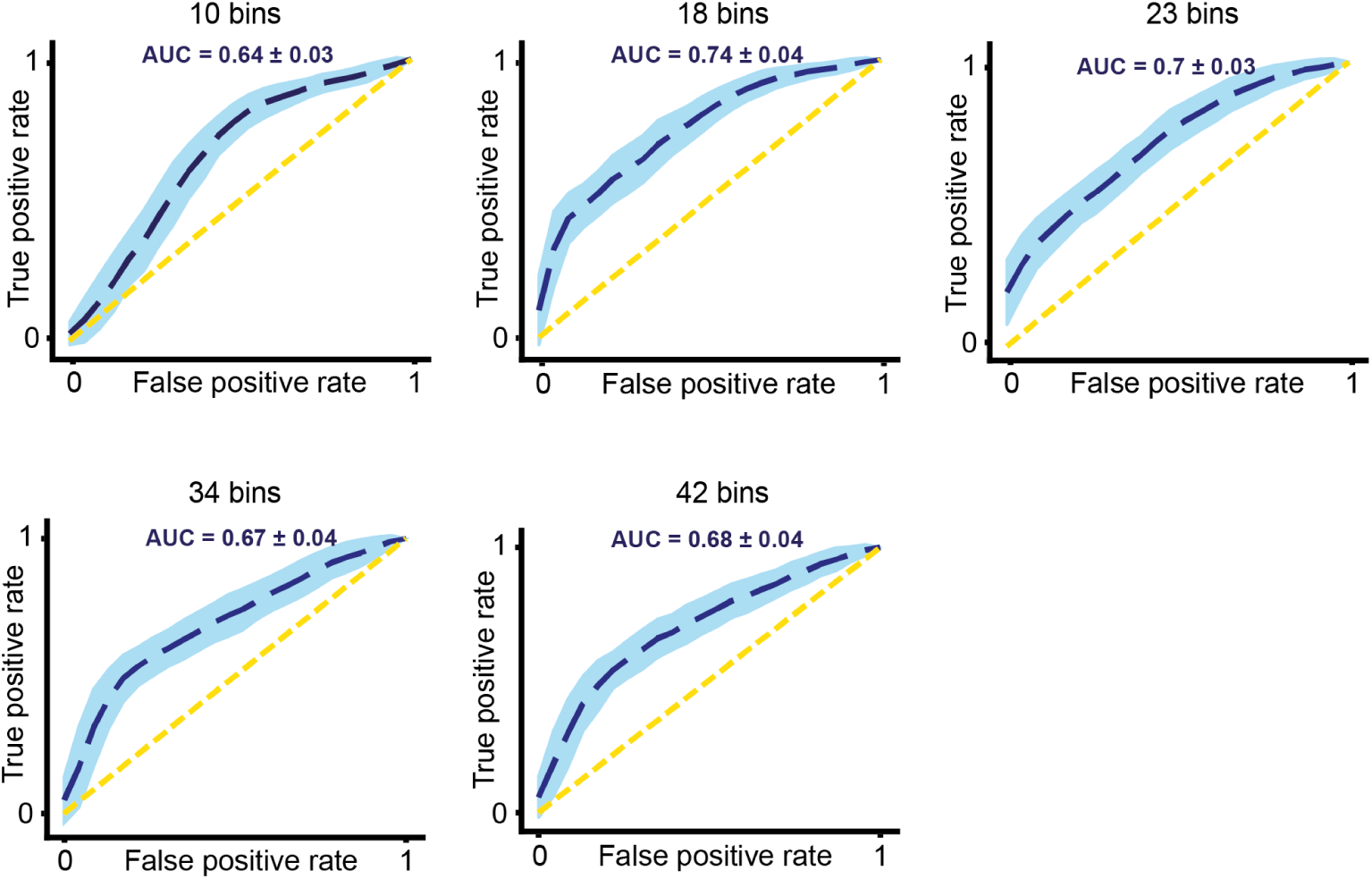
Receiver Operating Characteristic (ROC) curves for tissue-wise distribution of single nuclei lifetime XGBoost models trained using different binning resolutions. Each subplot shows the average ROC curve (blue dashed line) and standard deviation (blue shaded area) computed over 100 random seeds for a specific number of bins used to define the nuclei lifetime distribution. The AUC and its standard deviation are reported in each panel. The yellow dashed line indicates the performance of a random classifier.

**Figure S9.**
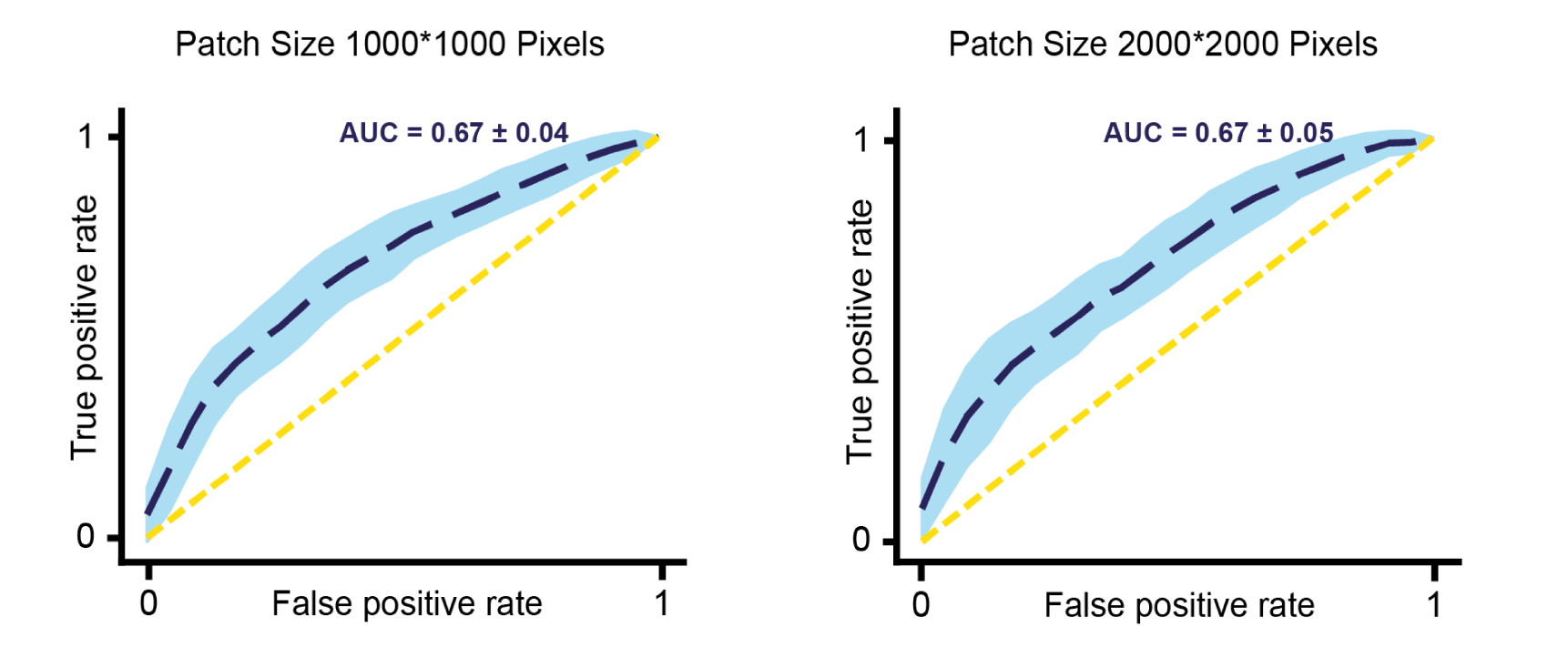
**Sensitivity of lifetime distribution patch-based prediction model to patch size**. ROC curves for models trained using lifetime distributions from patches of sized 1000×1000 pixels (left) and 2000×2000 pixels (right). The blue dashed lines represent the average ROC across 100 random seeds, and the shaded areas denote standard deviation. AUC values are reported in each panel. The yellow dashed lines indicate random classification.

**Figure S10.**
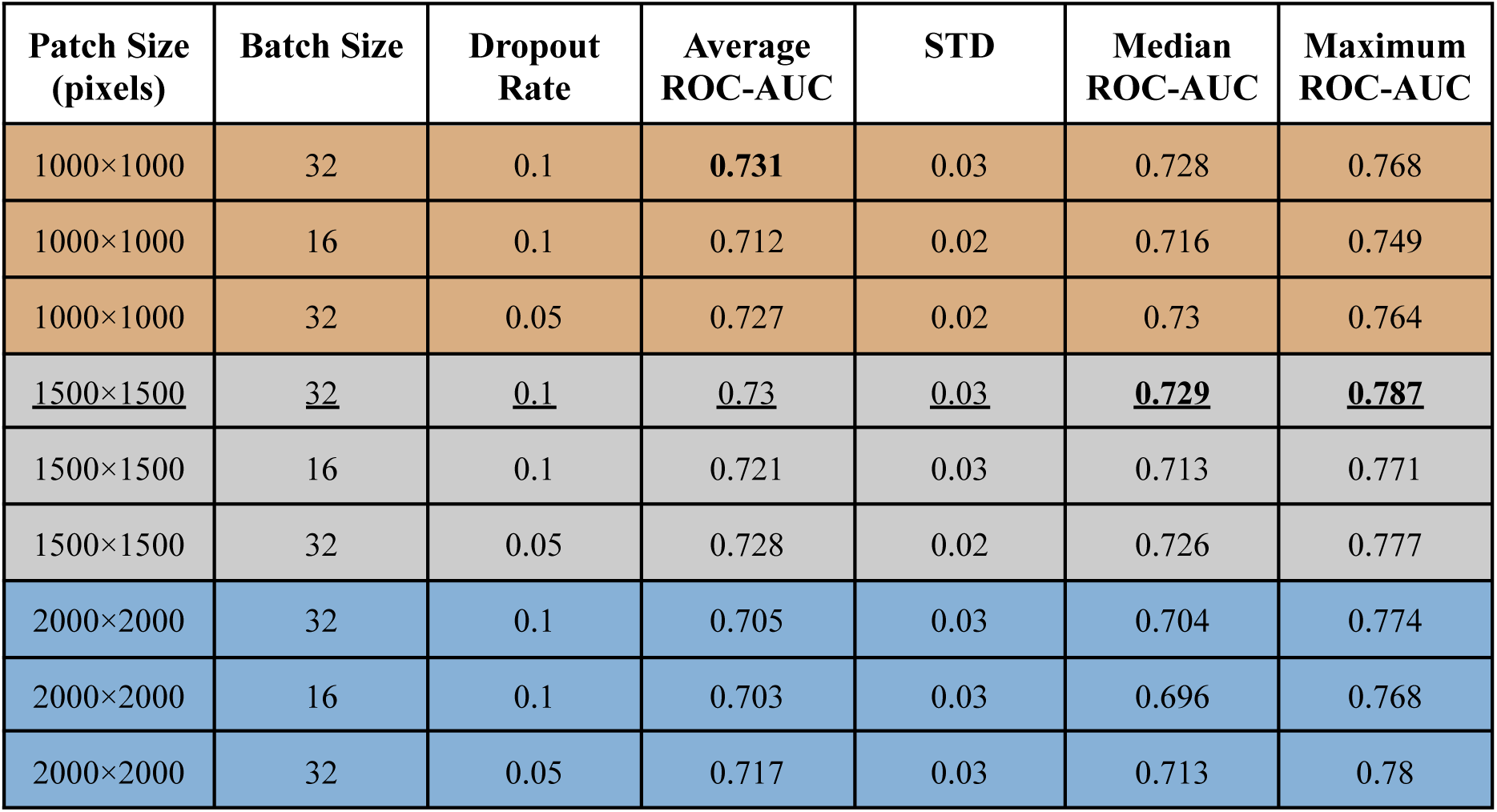
Hyperparameter sensitivity analysis of the GNN model to patch size. The performance of GNN models trained with different hyperparameters: patch size (1000×1000, 1500×1500, 2000×2000 pixels), batch size (16, 32), and dropout rate (0.1, 0.05). Metrics include average ROC-AUC, standard deviation (STD), median ROC-AUC, and maximum ROC-AUC, all computed across 20 random seeds. Bold corresponds to the best value in a metric and underlines represent the configuration we used.

